# Functional impact of PCSK9 variants on LDL uptake in a knockout hepatic model

**DOI:** 10.64898/2026.03.13.711724

**Authors:** Hongmei Li, Huili Liu, Wenxin Xu, Yuanjun Zeng, Pengfei Huang, Jiawang Guo, Binxiang Cai, Yajing Chen, Yanyan Lin, Chunbin Zhang

## Abstract

Proprotein convertase subtilisin/kexin type 9 (PCSK9) is a central regulator of low-density lipoprotein (LDL) cholesterol metabolism, yet the functional consequences of many clinically observed PCSK9 variants remain unknown. To establish a rigorous system for quantitative variant assessment, we generated a PCSK9 knockout (KO) HepG2 cell line through CRISPR/Cas9-mediated deletion of exons 2–8, effectively removing both the prodomain and catalytic regions required for PCSK9 function. This null background enabled systematic functional mapping of wild-type (WT) PCSK9 and multiple clinically relevant variants representing well-characterized, recurrent, and previously understudied alleles. Functional assays revealed pronounced heterogeneity among variant activities. The classical gain-of-function (GOF) variants D374Y and R496W exhibited robust suppression of LDL uptake, whereas A443T—an infrequently reported and previously uncharacterized variant—demonstrated a loss-of-function (LOF)-like phenotype with significantly enhanced LDL uptake. Additional poorly characterized variants, including V4I, R104C/V114A, and R496W/N425S, displayed minimal functional profiles, providing novel mechanistic insights. Surface LDL receptor (LDLR) levels generally correlated with LDL uptake but revealed unique patterns for specific variants. This KO-based rescue system provides a high-resolution framework for mechanistic classification of both established and poorly characterized PCSK9 variants, bridging the gap between genetic discovery and functional interpretation while supporting precision lipid-lowering strategies.

## Introduction

PCSK9 plays an indispensable role in cholesterol metabolism by binding to the low-density lipoprotein receptor (LDLR) and promoting its lysosomal degradation, thereby reducing hepatic LDL clearance. The clinical importance of this pathway has been underscored by the success of PCSK9 inhibitors, which significantly reduce LDL cholesterol (LDL-C) and cardiovascular events in high-risk patients(1–4). Concurrently, population and clinical sequencing studies have revealed extensive allelic diversity in PCSK9, including loss-of-function (LOF) variants associated with reduced LDL-C and cardiovascular protection(5–9), and gain-of-function (GOF) variants that contribute to familial hypercholesterolemia(10–13).

Despite this rapidly expanding genetic landscape, many PCSK9 variants remain functionally uncharacterized, limiting their clinical interpretability. Well-known variants such as D374Y and R46L have documented effects, yet numerous additional missense and compound variants—including A443T, V4I, R104C/V114A, and R496W/N425S—lack rigorous functional validation. This represents a growing challenge, as accurate variant interpretation is increasingly essential for precision lipid management and genetic counseling(14, 15).

A major limitation of current in vitro PCSK9 studies is the presence of endogenous PCSK9 in most cellular systems, which obscures subtle variant-specific effects. Prior investigations commonly relied on overexpression or partial knockdown approaches, restricting mechanistic resolution. To address these constraints, we engineered a large-deletion PCSK9 knockout HepG2 line that eliminates exons 2–8, including the prodomain and catalytic domain required for autocatalytic maturation and LDLR binding(16). This strategy allows reintroduction of defined PCSK9 variants into a PCSK9-null background under standardized expression conditions.

Using this platform, we systematically evaluated a panel of clinically relevant PCSK9 variants spanning well-studied pathogenic alleles and previously understudied or uncharacterized variants. By integrating LDL uptake and surface LDLR quantification, we delineate a comprehensive functional spectrum that highlights novel mechanistic properties of several rare variants. Our findings provide new insights into PCSK9 structure–function relationships and expand the functional annotation necessary for accurate variant interpretation in clinical genomics.

## Materials and Methods

### Cell Culture and Transfection

HepG2 hepatocellular carcinoma cells (HyCyte, Suzhou, China, Catalog No. TCH-C196) were selected as the experimental model due to their hepatic origin and endogenous expression of cholesterol metabolism machinery. Cells were cultured in complete MEM medium supplemented with 10% fetal bovine serum and 1% penicillin-streptomycin at 37°C in a humidified atmosphere with 5% CO₂.

For transfection experiments, cells were seeded in 6-well plates at a density of 4×10⁵ cells per well and allowed to adhere overnight. Transfection was performed using a lipofection-based approach with the following system: Solution A contained 100 μL Opti-MEM, 3 μg DNA, and 6 μL p3000 reagent; Solution B contained 100 μL Opti-MEM and 6 μL Lip3000 reagent. Both solutions were incubated separately at room temperature for 5 minutes, then combined and incubated for an additional 20 minutes before dropwise addition to cells. Cells were cultured for 48 hours post-transfection to allow sufficient protein expression before experimental analysis.

### Guide RNA Design

Single guide RNAs (sgRNAs) were designed using the CRISPOR web-based platform (http://crispor.tefor.net) for comprehensive analysis of potential targeting sites and off-target prediction (17). After systematic evaluation, the final selected gRNAs were gRNA-A1 (5’-GGTGAGATAAAGTACACCTA-3’) with PAM sequence GGG targeting a sequence upstream of exon 2, and gRNA-A2 (5’-ACATGCTGTCCACACAATGG-3’) with PAM sequence CGG targeting a sequence downstream of exon 8. These gRNAs were predicted to create a deletion of approximately 17.9 kb, removing exons 2-8 . Exons 2 through 8 were selected as the target region because this span encompasses critical functional domains including the prodomain (exons 2-3), which is essential for autocatalytic processing and protein maturation, and the catalytic domain (exons 4-8), which contains the serine protease active site and LDLR binding interface (16).

### Ribonucleoprotein (RNP) Complex Assembly and Electroporation

Prior to electroporation, HepG2 cell viability was assessed using trypan blue exclusion assay, with only cultures showing >95% viability being used for transfection. Cells were harvested during logarithmic growth phase (70-80% confluency) to ensure optimal electroporation efficiency and post-transfection recovery.

High-quality synthetic gRNAs were chemically synthesized with 2’-O-methyl 3’-phosphorothioate modifications to enhance stability and reduce immunogenicity (18). Ribonucleoprotein complex formation involved incubating each gRNA (final concentration 61 μM) with Cas9 protein (31 μM) in a 2:1 molar ratio for 10 minutes at room temperature. Electroporation was performed using the Neon™ Transfection System with optimized electrical parameters (1,350 V, 10 ms, 3 pulses). Cells were immediately transferred to pre-warmed recovery medium and incubated for 24 hours before analysis. EGFP mRNA electroporation served as a positive control to assess transfection efficiency.

### Single Cell Isolation and Molecular Screening

Following a 48-hour recovery period, cells were trypsinized and seeded at limiting dilution densities (0.5 cells/well) in 96-well plates containing conditioned medium. Wells containing single cells were identified by microscopic examination and monitored daily for colony formation. Individual clones were expanded over 2-3 weeks with regular medium changes, and once colonies reached approximately 100-200 cells, they were transferred to 24-well plates for further expansion.

A three-primer set strategy was designed to comprehensively characterize knockout events. Region 1 primers (forward: 5’-TCCCTTCTGCCTGCATTTGT-3’, reverse: 5’-CCCCATGCAAGGAGGAACAT-3’) were designed to amplify a 588 bp wild-type fragment that would be absent in homozygous knockouts. Region 2 primers (forward: 5’-TCAAGGAGCATGGAATCCCG-3’, reverse: 5’-TGCCTTGCTGGTCATGCTAA-3’) amplified a 611 bp wild-type fragment also absent in knockouts. Region 3 primers (forward: 5’-TCCCTTCTGCCTGCATTTGT-3’, reverse: 5’-GCCTTGCTGGTCATGCTAAC-3’) were designed to span the deletion junction, producing an 18,881 bp wild-type amplicon or a ∼970 bp knockout-specific amplicon.

PCR conditions were optimized with an annealing temperature of 62.0°C using high-fidelity polymerase. Initial screening involved standard PCR followed by agarose gel electrophoresis, with positive clones identified by the presence of the knockout-specific band and absence of wild-type bands. Confirmed knockout clones underwent Sanger sequencing for precise molecular characterization of deletion breakpoints.

### PCSK9 Variant Plasmid Construction and Transfection

Full-length human PCSK9 (NCBI RefSeq: NM_174936.4) and its variants were synthesized and subcloned into a pLVX-Puro-Myc vector. Variants selected in our study were those were those identified across multiple ethnic groups, including Chinese populations(11). All plasmid sequences were verified by Sanger sequencing. For overexpression experiments, HepG2 cells were seeded in 6-well plates and transfected using a lipofection-based reagent according to the manufacturer’s protocol. Cells were cultured for 48 hours post-transfection before analysis.

### Western Blot Analysis

Protein extraction was performed using enhanced RIPA lysis buffer containing protease inhibitors. Cells were washed three times with PBS, then lysed on ice for 20 minutes. Following homogenization and centrifugation at 12,000 rpm for 2 minutes at 4°C, supernatants were collected for protein quantification.

Protein concentrations were determined using the BCA assay with bovine serum albumin standards. Working solution was prepared by mixing reagents A and B at a 50:1 ratio. Standard protein concentrations were prepared in eight gradients (0, 1, 2, 4, 6, 8, 10, 20 μL), and samples were incubated at room temperature for 20 minutes before measuring absorbance at 562 nm. Protein samples were normalized to 20-60 μg per 20 μL and mixed with 5× loading buffer before denaturation at 100°C for 10 minutes.

SDS-PAGE was performed using 10% polyacrylamide gels with 5% stacking gel. Electrophoresis was conducted at 80V for 30 minutes, then 120V for 80 minutes until the bromophenol blue dye front reached 1 cm from the gel bottom. Proteins were transferred to nitrocellulose membranes at constant current (400 mA) for 50 minutes at 4°C. Membranes were blocked with 5% BSA in TBST for 1 hour at room temperature with gentle shaking.

Primary antibodies against PCSK9 (ab181142, Abcam) or GAPDH (10494-1-AP, Proteintech, Wuhan, China) were diluted in TBST (1:1000 and 1:5000, respectively) according to manufacturer specifications and incubated overnight at 4°C with gentle agitation. Following three 10-minute washes with TBST, membranes were incubated with appropriate secondary antibodies (SA00001-2 and SA00001-1 from Proteintech, 1:10000) for 1 hour at room temperature. After five additional 5-minute TBST washes, protein detection was performed using ECL chemiluminescent substrate (1:1 mixture of solutions A and B) with 1-minute incubation before imaging and analysis.

### RNA Extraction and Quality Assessment

Total RNA was extracted using RNAiso Plus reagent following a modified phenol-chloroform protocol optimized for high yield and purity (19). RNA concentration and purity were assessed using nucleic acid analyzer measuring absorbance at 260 nm and 280 nm. RNA extraction from all transfected cell populations yielded consistently high-quality samples suitable for downstream molecular analysis. Concentrations ranged from 176.8 ng/μL to 870.9 ng/μL, with A260/A280 ratios consistently falling between 1.809 and 1.919 across all samples, well within the acceptable range indicating minimal protein contamination and excellent RNA purity. First-strand cDNA synthesis was performed using 1 μg total RNA, followed by quantitative PCR using SYBR Green chemistry with gene-specific primers designed to span exon-exon junctions (20).

### Quantitative PCR Analysis

First-strand cDNA synthesis was performed using 1 μg total RNA with NovoScript® 1st Strand cDNA Synthesis SuperMix. The reaction was incubated at 50°C for 15 minutes, followed by 85°C for 5 seconds to inactivate the reverse transcriptase.

Real-time PCR was conducted using SYBR Green chemistry with gene-specific primers designed using Primer5 software and validated by BLAST analysis. PCSK9 primers were: forward 5’-AGACCCACCTCTCGCAGTC-3’, reverse 5’-GGAGTCCTCCTCGATGTAGTC-3’. GAPDH served as the internal reference gene with primers: forward 5’-GAAGGTCGGTGTGAACGGAT-3’, reverse 5’-CCCATTTGATGTTAGCGGGAT-3’.

The 20 μL reaction mixture contained 10 μL 2× SYBR qPCR Mix, 0.5 μL each of forward and reverse primers (10 μM), 1 μL cDNA template, and 8 μL nuclease-free water. Thermal cycling conditions included initial denaturation at 95°C for 10 minutes, followed by 40 cycles of 95°C for 15 seconds and 60°C for 30 seconds. Melting curve analysis was performed from 60°C to 95°C to verify primer specificity. Relative expression was calculated using the 2-ΔΔCt method with GAPDH normalization.

### LDL Uptake Functional Assay

Cellular LDL uptake was measured using fluorescently labeled 1,1’-dioctadecyl-3,3,3’,3’-tetramethylindocarbocyanine-labeled acetylated low-density lipoprotein (DiI-Ac-LDL), which provides quantitative assessment of functional LDL receptor activity (21). Following transfection and appropriate recovery periods, cells were incubated with DiI-Ac-LDL diluted to 40 μg/mL in serum-free culture medium for 4 hours at 37°C.

After incubation, cells were washed three times with probe-free medium to remove unbound LDL particles and minimize background fluorescence. LDL uptake was quantified by measuring the fluorescence intensity of internalized DiI-Ac-LDL particles using flow cytometry. Data were collected from at least 10,000 cells per sample to ensure statistical reliability.

### Quantitation of Cell Surface LDLR Receptor

HepG2 cells were seeded at 4×10^5^ cells per well in 6-well plates and allowed to reach confluence before transfection. Transient transfections were performed using Lipofectamine 3000 reagent according to manufacturer’s protocol, with 3 μg DNA per well and a 20-minute room temperature incubation period. Experimental groups included parent HepG2 cells, PCSK9 knockout (KO) control clone 1G9, 1G9 overexpressing wild-type PCSK9 (WT) and various PCSK9 variants (D374Y, R496W, E32K, V4I, R46L, R104C/V114A, R496W/N425S, and A443T). Following 48-hour transfection, cells were dislodged, washed twice with PBS, and processed for flow cytometric analysis. For LDL receptor (LDLR) detection, 10^6^ cells were blocked with 1% FBS for 30 minutes at room temperature, incubated with recombinant APC anti-LDLR antibody (Abcam, ab275614) for 1 hour at room temperature in the dark, washed twice with PBS, and resuspended in 200 μL PBS for flow cytometric analysis using an Agilent NovoCyte 2060R flow cytometer. Data were analyzed by Flow Jo 10.8.1.

### Statistical Analysis

All quantitative data were analyzed using GraphPad Prism 8 software with appropriate statistical tests selected based on data distribution and experimental design. Statistical comparisons between experimental groups were performed using one-way analysis of variance (ANOVA) with Tukey’s multiple comparisons test when comparing multiple groups, or unpaired t-tests for pairwise comparisons. Results are presented as mean ± standard error of the mean (SEM) from at least three independent experiments, with statistical significance levels set at P < 0.05.

## Results

To comprehensively investigate the role of PCSK9 role in cholesterol metabolism, we employed a systematic experimental approach combining both loss-of-function and gain-of-function strategies. This dual methodology provides complementary evidence while enabling detailed functional analysis of multiple mutant variants. We chose HepG2 cells in the study because of their hepatic origin and robust expression of cholesterol metabolism machinery.

### Generation and Molecular Validation of PCSK9 Knockout Cell Line

The human PCSK9 gene (Gene ID: 255738) was comprehensively analyzed using NCBI and Ensembl databases to determine optimal targeting strategies for generating knockout cell lines. PCSK9 is located on chromosome 1q32.1 and contains 12 exons spanning approximately 25 kb of genomic DNA (19). We designed two gRNAs to create a deletion of approximately 17.9 kb, removing exons 3-7 entirely and portions of exons 2 and 8 (Fig. 1A). Exons 2 through 8 were selected as the target region because this span encompasses critical functional domains including the prodomain (exons 2-3), which is essential for autocatalytic processing and protein maturation, and the catalytic domain (exons 4-8), which contains the serine protease active site and LDLR binding interface(16) (Fig. 1B). Of note, natural amino acid changes, resulting in different PCSK9 variants, have been reported to various PCSK9 domains with clinical implications (Fig. 1C).

**Fig. 1.**
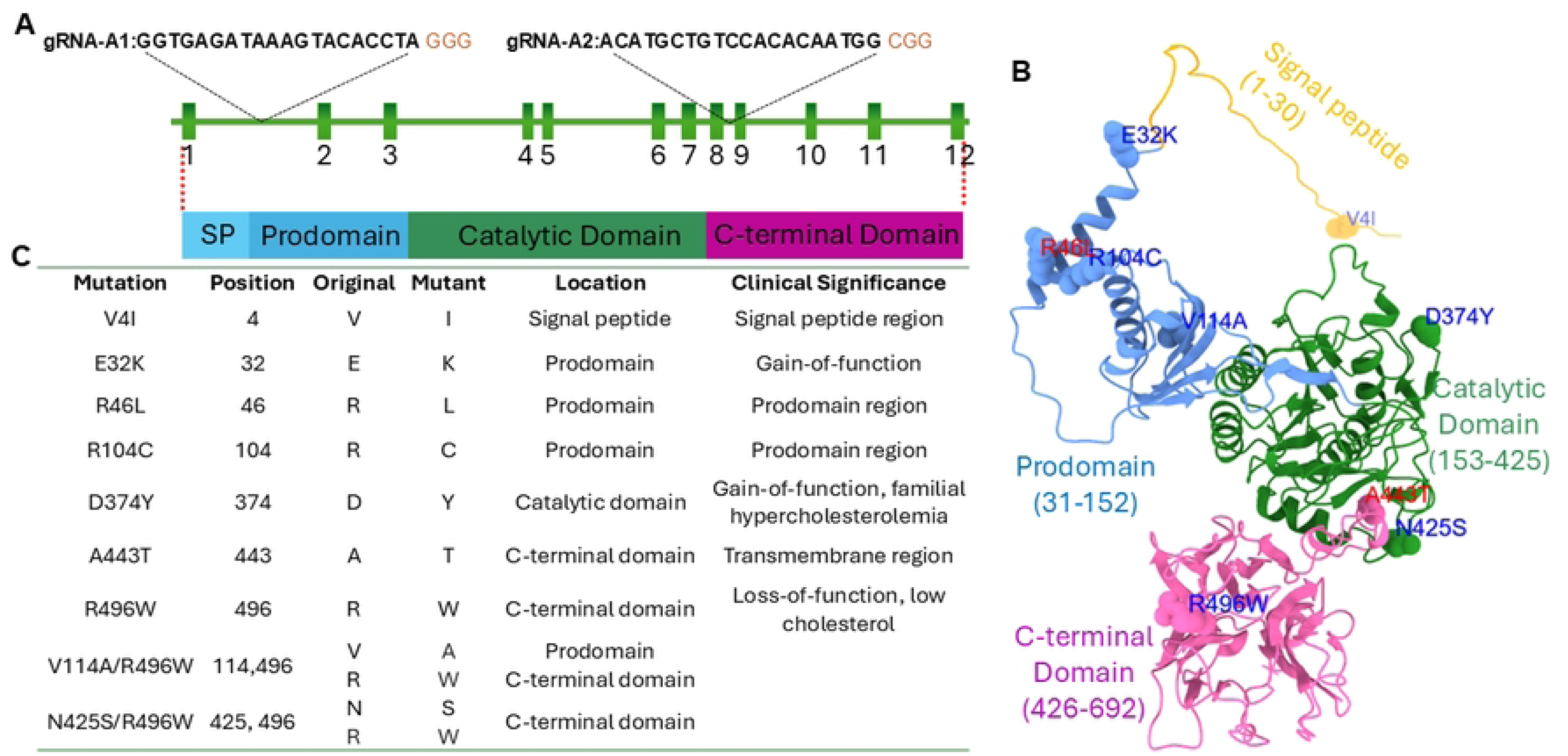
Human PCSK9 and variants. (A) Schematic of the human PCSK9 gene on chromosome 1, indicating the 12 exons. The positions of the two guide RNAs (gRNAs) targeting exon 2 and exon 8 are shown. A predicted ∼17.9 kb deletion will remove exons 2-8. This region encompasses the prodomain and the catalytic domain. (B) Diagram of the PCSK9 protein domains (Prodomain, Catalytic Domain, C-terminal Domain) with the locations of the clinically relevant amino acid variants investigated in this study. (C) A summary of variants to be investigated in this study.

We subsequently assembled gRNA-Cas9 Ribonucleoprotein (RNP) Complex and electroporated HepG2 cells with PCSK9-targeting RNP complexes. A comprehensive three-primer PCR strategy was designed to systematically characterize knockout events and distinguish between homozygous knockouts, heterozygous modifications, and wild-type cells (Details are described in Methods). Clone 1E4 and 1G9 demonstrated the desired genotype profile consistent with homozygous knockout and were selected for detailed characterization (Fig. 2).

**Fig. 2.**
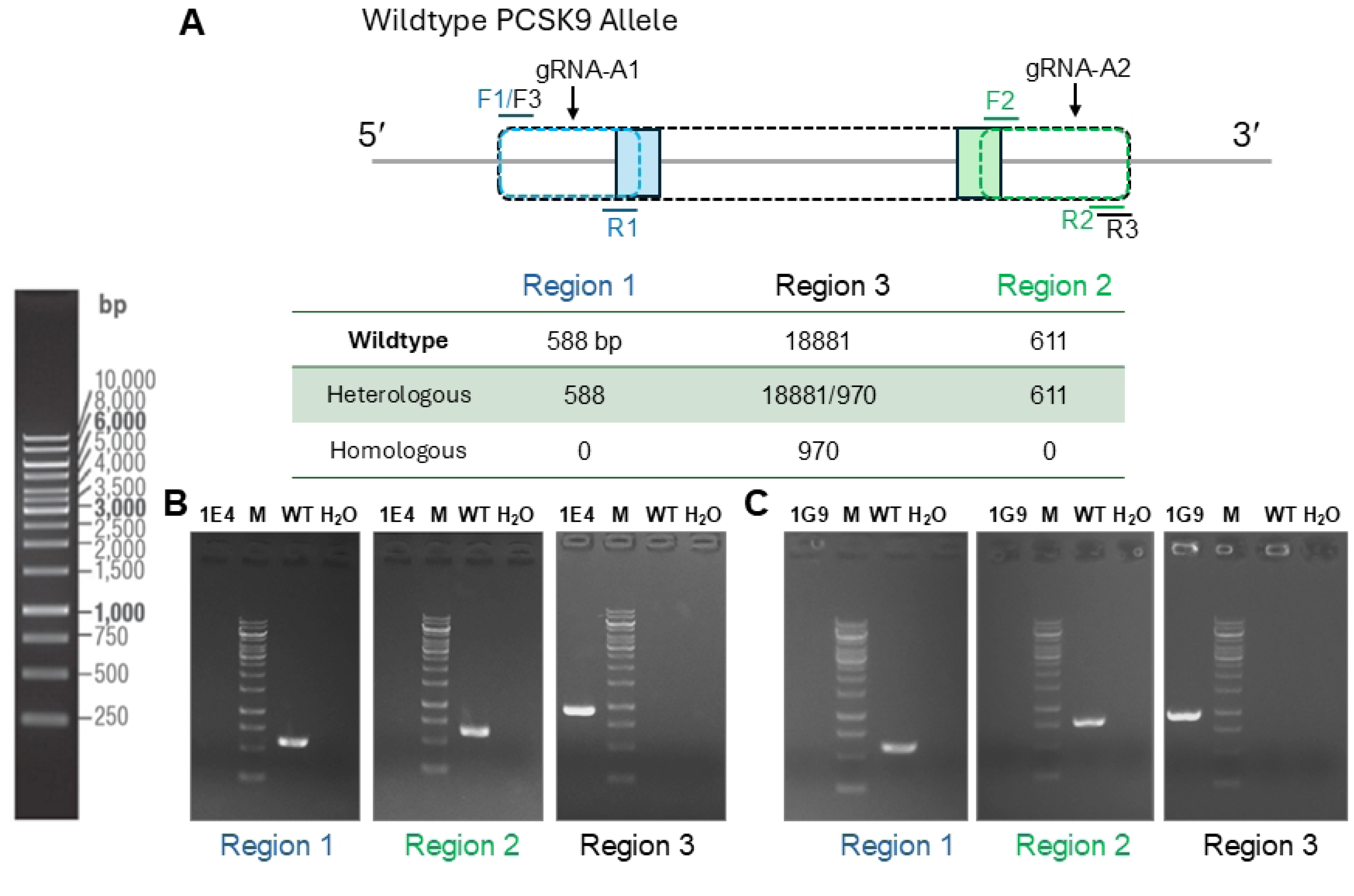
Generation of PCSK9 Knockout HepG2 Cell Clones. (A), Schematic of the CRISPR/Cas9 targeting strategy showing gRNAs targeting exons 2 and 8 of the PCSK9 gene. Comprehensive PCR screening of the clones using the three-primer strategy reveals homozygous or heterozygous clones. Region 1 and Region 2 primers amplify WT-specific fragments that are absent in both clones. Region 3 primers, designed to span the deletion, amplify a large ∼18.9 kb band in WT cells and a distinct ∼970 bp knockout-specific band in clones 1E4 and 1G9, confirming a homozygous deletion event. (B), Agarose gel electrophoresis of PCR products from the three-primer screening strategy for wild-type (WT) HepG2 cells and two representative single-cell clones (1E4 and 1G9). M: DNA ladder.

Growth characteristics analysis showed that both clones maintained normal HepG2 morphology with epithelial-like appearance and contact inhibition, with a doubling time of 28 ± 4 hours comparable to parental cells (26 ± 3 hours) (Fig. 3A), but western blotting revealed that there is residual PCSK9 protein in Clone 1E4. By contrast, a complete ablation of PCSK9 protein expression was observed in clone 1G9 compared to wild-type controls, with the characteristic ∼ 62 kDa PCSK9 band clearly visible in wild-type HepG2 cells but completely absent in 1G9 cells (Fig. 3B). This clean knockout profile confirmed successful gene disruption without detectable compensatory proteins or truncated gene products (22). To confirm the genetic knockout of PCSK9, genomic DNA isolated from 1E4 and 1G9 cells were subjected to Sanger sequencing. Clone 1E4 contains a precise 17,930 base pair deletion between the two gRNA target sites, encompassing targeted exonic regions. Sanger sequencing of clone 1G9 revealed a large deletion of 17,920 bp spanning from the intron 1/exon 2 boundary to the intron 7/exon 8 boundary. A secondary sequencing analysis revealed a slightly different deletion size (17,911 bp), suggesting minor variability in the exact breakpoint positions, which is characteristic of CRISPR-mediated deletions due to non-homologous end joining repair mechanisms(23) (Fig. 3C).

**Fig. 3.**
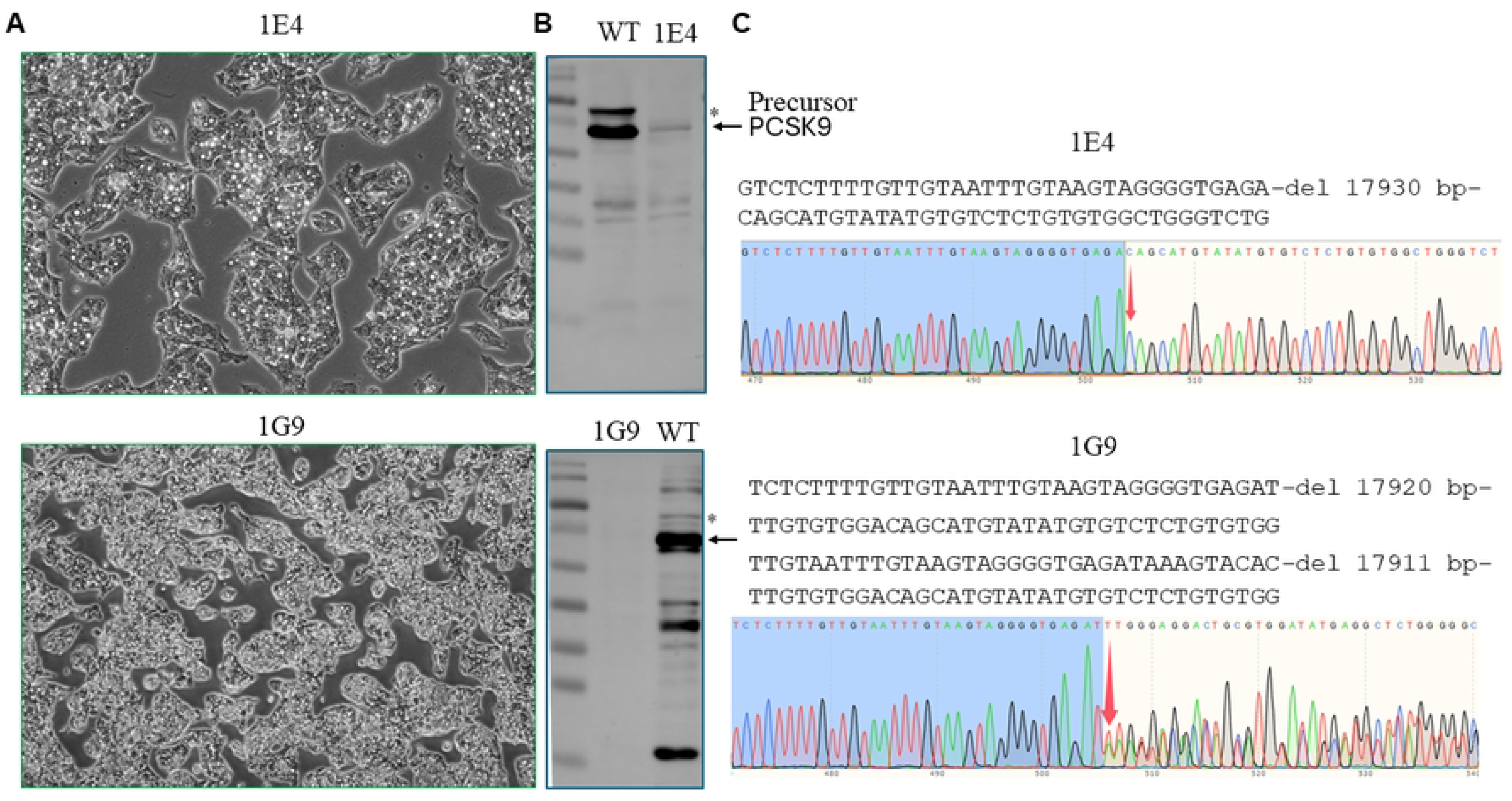
Validation of the PCSK9 Knockout (KO) Clone 1G9. (A) Normal epithelial-like morphology and contact inhibition for both 1E4 and 1G9 KO clones. Images were taken under 100x and 40x objectives using an Olympus SZX7 stereo microscope. (B) Western blot analysis of PCSK9 protein expression. The characteristic 62 kDa PCSK9 band is present in wild-type (WT) cells but completely absent in the 1G9 KO clone. The * indicates PCSK9 precursor (Pro-PCSK9), which migrates to a position higher than 70 kDa. The mature form of PCSK9 migrates around 62 kd. Clone 1E4 shows residual PCSK9 expression. Th protein ladder is Thermo Scientific PageRuler Prestained Protein Ladder (#26616). (C) Sanger sequencing chromatogram of the PCR product from clone 1G9, confirming a precise 17,920 bp genomic deletion between the gRNA target sites.

### Genetic Knockout of PCSK9 Enhances Cellular Uptake of LDL and upregulates LDLR

Using clone 1G9, we subsequently evaluated the consequences of PCSK9 deficiency on cellular cholesterol uptake capacity. To this end, 1,1’-dioctadecyl-3,3,3’,3’-tetramethylindocarbocyanine-labeled acetylated low-density lipoprotein (DiI-Ac-LDL) was added to parent HepG2 cells and the PCSK9 KO 1G9 clone. Flow cytometric analysis of uptake of fluorescently labeled DiI-Ac-LDL clearly demonstrated that 1G9 cells exhibited significantly increased fluorescence intensity, indicating substantially greater internalization of DiI-Ac-LDL particles (Fig. 4A&B). Concomitantly, the cell surface level of LDL receptor (LDLR) also slightly increased in 1G9 cells (Fig. 4C). These findings provide direct functional evidence that PCSK9 negatively regulates LDL uptake, consistent with its established role in promoting LDL receptor degradation and reducing hepatic cholesterol clearance capacity (24).

**Fig. 4.**
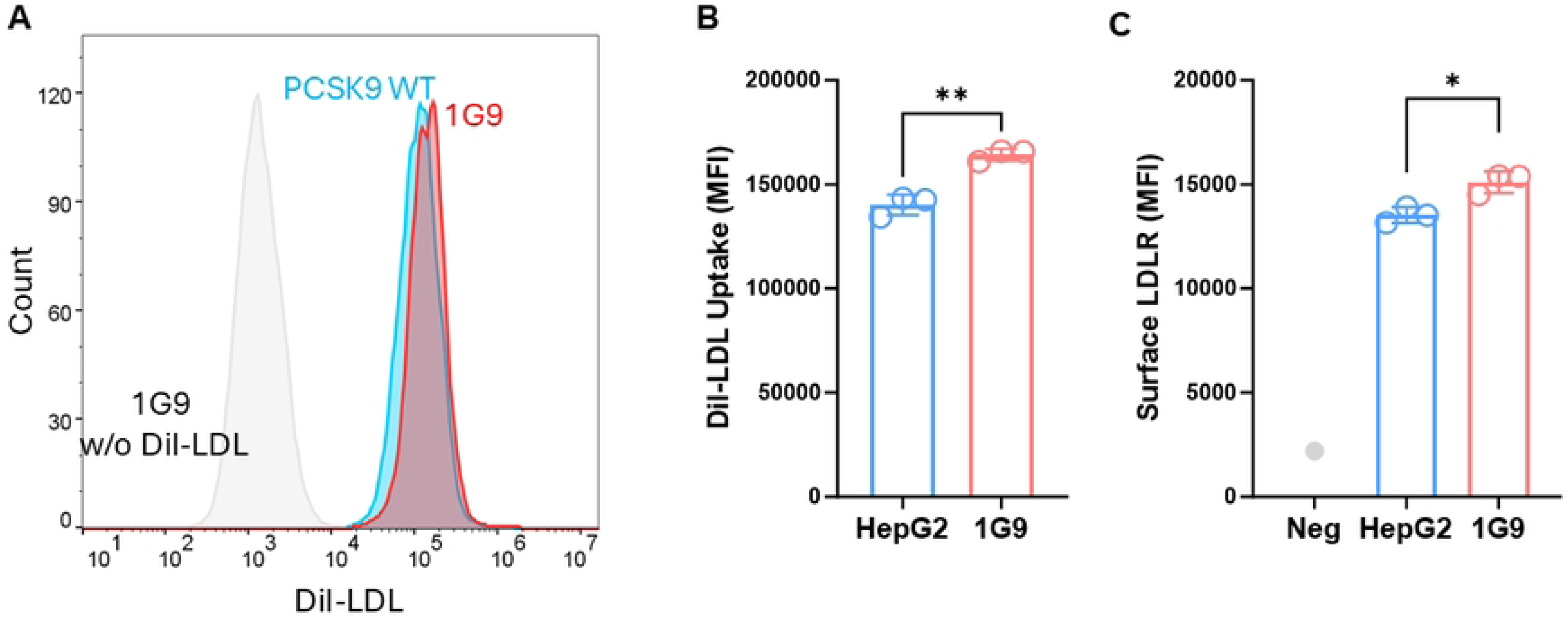
Functional Validation of PCSK9 Knockout via LDL Uptake Assay. (A) Flow cytometry analysis of DiI-Ac-LDL uptake in wild-type (WT) and PCSK9 knockout (KO) HepG2 cells. The histogram shows a rightward shift in fluorescence intensity for the KO clone (red) compared to WT cells (blue). (B) The mean fluorescence intensity of DiI-Ac-LDL was also presented. This result provides functional confirmation that loss of PCSK9 increases cellular LDL uptake. **p<0.005. (C) Surface expression of LDLR on HepG2 and 1G9 cells was quantified by flow cytometry. *p<0.05.

### Varied Impacts of PCSK9 Variants on Cellular Uptake of LDL

Having established the baseline importance of PCSK9, we next developed overexpression systems to explore direct impacts of PCSK9 mutants on cellular LDL uptake. To this end, 1G9 cells were transfected with DNA plasmids expressing wild-type PCSK9 (WT), multiple variants with single amino acid change (D374Y, R496W, E32K, V4I, R46L, A443T), and two compound variants (R104C/V114A, R496W/N425S), providing broad coverage of different mutation types and their potential effects. Quantitative PCR analysis confirmed overexpression of PCSK9 variants in transfected cells to comparable levels (Fig. 5A). In contrast to the RNA-level validation, Western blot analysis revealed the expression of PCSK9 variants at the protein level varied significantly, with multiple protein specifies noted (Fig. 5B&C), suggesting that some amino acid changes may affect protein stability, post-translational modifications, or electrophoretic mobility (25). For example, PCSK9 variant containing R104C/V114A was expressed to a much lower level, consistent with the previously reported phenotype of impaired processing in hepatocyte cell cultures (26).

**Fig. 5.**
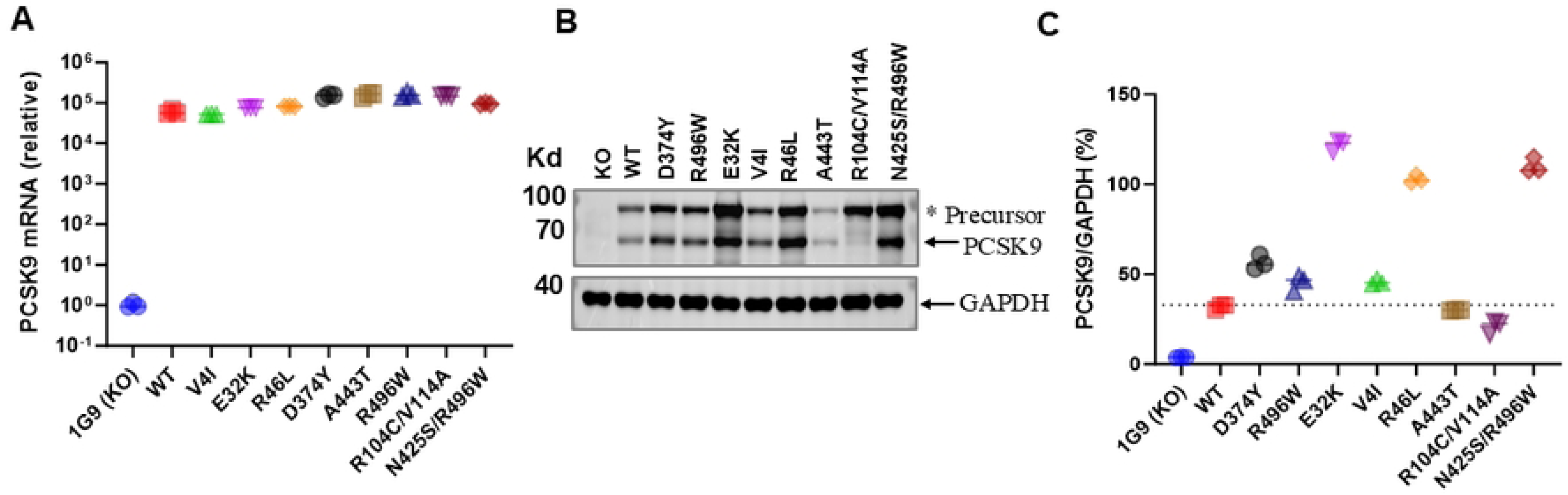
Validation of PCSK9 Variants Overexpression in 1G9 Cells. (A) qPCR amplification for PCSK9 and then normalized to GAPDH reference gene is presented to confirm specific and robust transcript expression of PCSK9 variants in transfected cells. (B) Western blot analysis showing expression of wild-type (WT) PCSK9 and all tested variants at the expected molecular weight of ∼62 kDa in transfected 1G9 KO cells. (C) Densitometric quantification of the Western blot bands, showing variable protein expression levels among the different variants.

### Understudied Variants Exhibit Previously Unrecognized Functional Properties

Quantitative LDL uptake assays revealed distinct functional signatures across variants (Fig. 6A). A443T, a rarely reported variant lacking prior mechanistic characterization, significantly increased LDL uptake relative to WT, identifying it as a LOF-like allele. The compound variant R104C/V114A produced modest increases in LDL uptake and surface LDLR levels (Fig. 6B), extending earlier reports of its processing defect by demonstrating functional consequences at the receptor level(26). The V4I variant displayed minimal deviation from WT for both LDL uptake and LDLR expression, suggesting functional neutrality. The compound variant R496W/N425S exhibited a phenotype like the WT, suggesting an intramolecular interaction that modifies the classical GOF effect exerted by R496W. Classical GOF variants D374Y and R496W showed strong suppression of LDL uptake, confirming assay fidelity. The R46L variant yielded only minor effects in this system, potentially reflecting intrinsic limitations of transient expression systems.

**Fig. 6.**
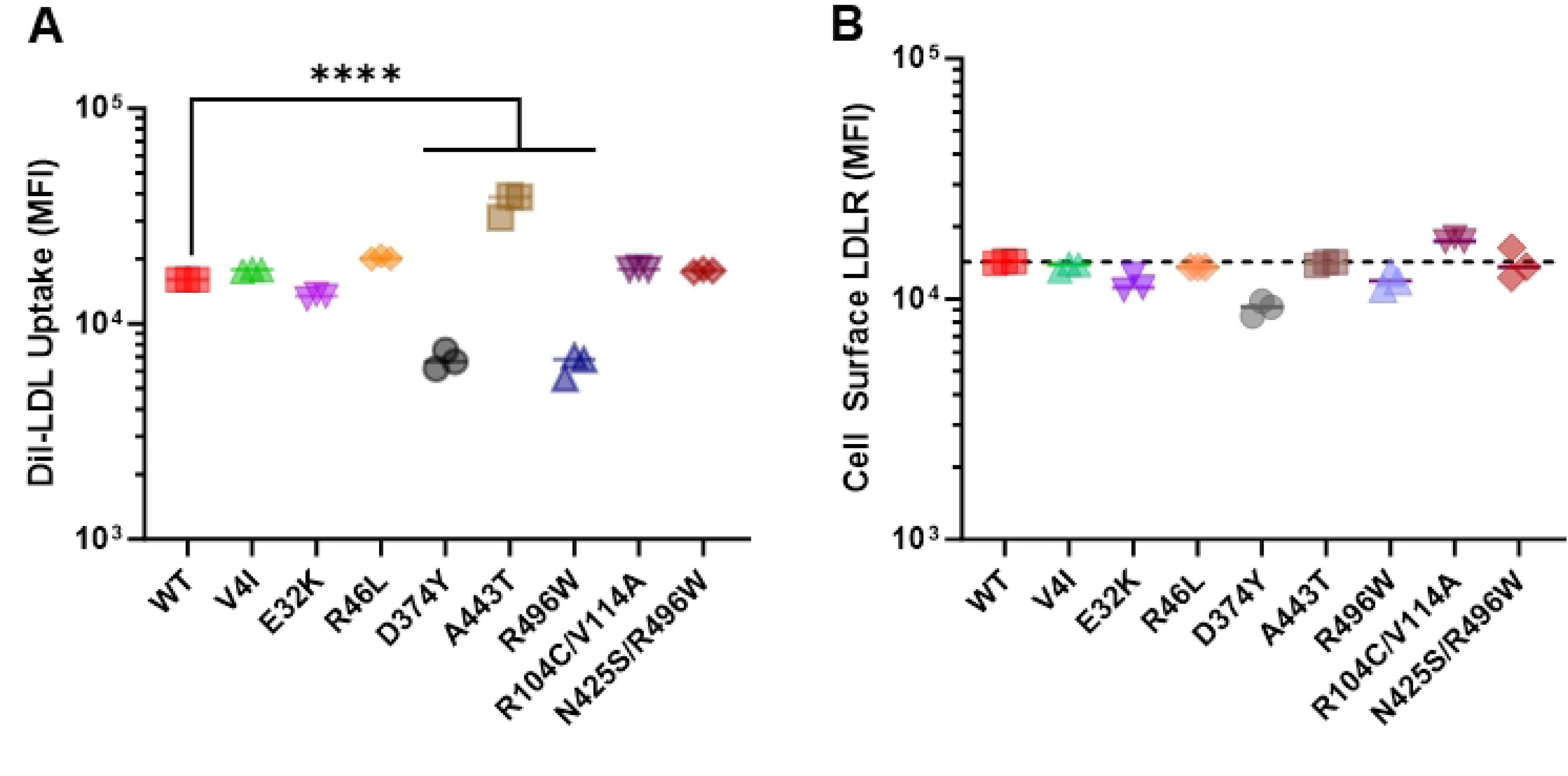
Functional Spectrum of PCSK9 Variants on LDL Uptake and LDLR Surface Expression. PCSK9 knockout (1G9) cells were transfected with a vector expressing wild-type (WT) PCSK9 or the indicated PCSK9 variants. (A) Quantification of DiI-Ac-LDL uptake by flow cytometry. Data were adjusted for the normalized expression level of each variant relative to the WT control. (B) Quantification of cell surface LDLR levels by flow cytometry. Data are presented as mean ± SEM from three independent experiments. Statistical significance was determined by one-way ANOVA test. (P < 0.05, *P < 0.01, *P < 0.001 compared to WT).

To investigate if the above findings correlate with the changes of LDL receptor (LDLR), which is known to be regulated by PCSK9, we quantified the cell surface LDLR in 1G9 cells overexpressing wild type or various variants. Compared to wild-type PCSK9, E32K, D374Y, and R496W slightly decreased the amount of cell surface LDLR. The variant containing R104C/V114A modestly enhanced LDLR on cell surface (Fig. 6B). In general, the effect of a specific variant on LDL uptake positively correlates with its effect on LDLR expression, but such a correlation is not always observed. For example, A443T increased cellular uptake of LDL, but did not significantly increase LDLR on the cell surface. It is possible that in our transient expression system, the effect of overexpressing a variant on surface LDLR level is simply not optimal given the degradation of LDLR by PCSK9 takes time. A persistent expression, i.e., stable expression, may result in more quantifiable changes of cell surface LDLR.

## Discussion

This study introduces a robust KO–rescue platform for precise functional assessment of PCSK9 variants and demonstrates its utility in characterizing both well-established and previously understudied alleles. The large-deletion PCSK9 KO line eliminates confounding endogenous activity and provides a controlled background in which variant-specific effects can be quantified with high resolution.

Our work validated important properties for several rare PCSK9 variants. A443T was partially functionally characterized as a loss-of-function variant(27). Likewise, the compound variant R104C/V114A, previously shown to be poorly processed and secreted(26), demonstrated measurable effects on LDL uptake and LDLR surface abundance. V4I appeared functionally neutral, while the compound variant R496W/N425S showed intermediate effect not predicted by either individual substitution. These findings align with some previous clinical observations that individuals with naturally occurring PCSK9 loss-of-function mutations exhibit significantly reduced plasma cholesterol levels and substantially decreased cardiovascular disease risk (8, 28). The gain-of-function variant D374Y represents a particularly significant finding with immediate clinical implications for cardiovascular medicine. The enhanced ability of D374Y to suppress LDL uptake provides a potential mechanistic explanation for certain forms of familial hypercholesterolemia that may not be explained by traditional LDL receptor mutations or other known genetic causes (29). Patients carrying such gain-of-function PCSK9 variants would be expected to exhibit elevated plasma cholesterol levels and increased cardiovascular risk through enhanced LDLR degradation, making them prime candidates for PCSK9-targeted therapies. The D374Y gain-of-function variant, located in the catalytic domain near the LDLR binding interface, likely enhances protein-protein interactions through the introduction of aromatic interactions and elimination of negative charge. This structural modification may increase binding affinity to LDLR or improve the efficiency of receptor trafficking to degradation pathways (30). Interestingly, the A443T variant showed enhanced LDL uptake, suggesting that this amino acid substitution in the C-terminal domain may attenuate PCSK9-mediated LDLR degradation and thereby increase functional receptor activity. However, this increase in LDL uptake was not accompanied by a proportional rise in steady-state surface LDLR levels. This apparent discrepancy indicates that enhanced LDL uptake may not be explained solely by increased receptor abundance at the plasma membrane. Because LDL uptake is a dynamic process influenced by ligand binding, internalization efficiency, endosomal trafficking, and recycling kinetics, it is possible that A443T alters receptor turnover or functional activity rather than simply elevating steady-state surface expression. Notably, our surface LDLR measurements reflect receptor abundance at a single time point, whereas the uptake assay integrates cumulative receptor function over time. Therefore, the current data do not distinguish between changes in receptor abundance and alterations in receptor kinetics or ligand-binding properties. Future studies employing pulse–chase approaches, receptor recycling assays, or binding affinity analyses will be required to delineate the precise mechanism underlying this phenotype.

Of note, our results are not always consistent with what was reported in the literature. The R46L substitution, occurring near the autocatalytic cleavage site in the prodomain, was previously reported to be a loss-of-function change that is linked to lower LDL cholesterol (9). In our study, however, R46L did not have significant effect cellular LDL uptake, nor did it significantly modulate the surface level of LDLR. Such discrepancy may be attributable to the experimental system being used in our study or limited modulatory effect exerted by R46L. The transient overexpression-based assay in this study may partially compress dynamic range and reduce sensitivity to subtle reductions in PCSK9 activity. Under these conditions, moderate impairments in PCSK9 function—such as those attributed to R46L—may appear attenuated compared to the effect size observed clinically. The finding that R496W/N425S counterbalances the GOF R496W is interesting and warrants further discussion. N425S seems to have been associated with both increased and reduced LDL cholesterol, and the variant has been functionally studied previously (31).

Nonetheless, our findings highlight the importance of experimental validation for variants of uncertain significance (VUS), particularly as clinical sequencing increasingly identifies rare alleles lacking functional annotation. Our platform enables scalable mechanistic evaluation and can be readily extended to additional PCSK9 variants or other genes involved in cholesterol metabolism (32).

Limitations include the use of HepG2 cells, which may not fully reproduce primary hepatocyte physiology(33), and the transient expression system, which may underrepresent long-term effects. Notably, DiI-Ac-LDL can be internalized by scavenger receptors in addition to, or preferentially over, classical LDLR in certain cellular contexts. Native DiI-LDL would provide a more LDLR-selective assessment. In our study, DiI-Ac-LDL uptake was used as a functional readout of cellular lipoprotein uptake capacity under conditions of PCSK9 modulation, rather than as a strictly LDLR-specific assay. Additionally, PCSK9-mediated LDLR degradation can occur through both intracellular interactions in the secretory pathway and extracellular binding to cell surface LDLR. The present overexpression system captures total cell-autonomous PCSK9 activity but does not distinguish between these mechanisms. Thus, while the assay enables comparative functional assessment across variants, it does not isolate extracellular effects that may predominate under physiological conditions. Last but not the least, gain-of-function phenotypes may arise through multiple mechanisms, including altered LDLR binding affinity, enhanced receptor internalization, increased protein stability, altered secretion efficiency, resistance to inhibitory interactions (e.g., with LDL particles), or changes in plasma half-life. The present assay measures net LDLR degradation under defined cellular conditions and does not distinguish among these mechanistic possibilities. Despite these limitations, strong concordance between observed phenotypes and known variant behaviors supports the validity of this approach. Overall, this study expands functional understanding of PCSK9 variants, clarifies mechanisms for several previously uncharacterized alleles, and provides a framework for integrating functional genomics into precision lipid management.

## Abbreviations

DiI-Ac-LDL: 1,1’-dioctadecyl-3,3,3’,3’-tetramethylindocarbocyanine-labeled acetylated low-density lipoprotein
GOF: gain-of-function
LDLR: low-density lipoprotein receptor
LOF: loss-of-function
PCSK9: proprotein convertase subtilisin/kexin type 9
RNP: ribonucleoprotein.

